# Identification of Human Transferrin Receptor as an Entry Co-receptor for Parvovirus B19 Infection of Human Erythroid Progenitor Cells

**DOI:** 10.64898/2026.04.02.715920

**Authors:** Shane McFarlin, Kang Ning, Xiujuan Zhang, Cagla Aksu Kuz, Wei Zou, Fang Cheng, Steve Kleiboeker, Mario Mietzsch, Jianming Qiu

## Abstract

Parvovirus B19 (B19V), a member of the genus *Erythroparvovirus* within the *Parvoviridae* family, infects human erythroid progenitor cells (EPCs) of bone marrow and fetal liver, and causes various hematological disorders. The minor capsid protein VP1 of B19V contains a unique N-terminal region (VP1u) that facilitates virus binding and internalization into EPCs via its receptor-binding domain (RBD). We previously identified tyrosine protein kinase receptor UFO (AXL) as a proteinaceous receptor for B19V infection of EPCs. In this study, we employed an ascorbate peroxidase 2 (APEX2)-based proximity labeling method to identify host proteins that are associated with B19V VP1u during entry. This analysis revealed human transferrin receptor 1 (hTfR) as a key host protein associated with VP1u. hTfR knockdown in UT7/Epo-S1 cells, a B19V-permissive human megakaryoblastoid leukemia cell line, showed significantly reduced B19V internalization and replication. Biolayer interferometry (BLI) assays confirmed a direct interaction between B19V VP1u and hTfR extracellular domain (ECD). Inhibition of VP1u interaction with hTfR ECD, either by a monoclonal antibody targeting the apical domain of the ECD or human ferritin, a natural ligand of hTfR that binds the apical domain, significantly reduced VP1u binding to hTfR, as well as B19V internalization and B19V replication in ex vivo-expanded EPCs. Furthermore, mutant RBD proteins that bear amino acid substitutions in the three helical domains nearly abolished RBD binding to hTfR and significantly reduced the ability to inhibit B19V infection of EPCs. Collectively, our findings establish hTfR as a B19V entry co-receptor that mediates B19V internalization into its natural host EPCs.

**Significance:** B19V causes severe hematological disorders, including transient aplastic crisis, chronic pure red cell aplasia, and hydrops fetalis, by selectively infecting erythroid progenitor cells (EPCs). Despite its clinical impact, no approved antivirals or vaccines exist, largely due to limited understanding of viral entry mechanisms. A unique feature of B19V is the externalization of the VP1 unique region (VP1u) from the viral capsid, which mediates receptor engagement. Our prior studies identified AXL as an attachment receptor for B19V. Here, we identify that human transferrin receptor 1 (hTfR) acts as a critical co-receptor that directly binds VP1u and promotes viral internalization. Inhibition of the VP1u-hTfR interaction by competitive binding of hTfR with either an anti-hTfR monoclonal antibody or human ferritin significantly reduces B19V internalization and replication in ex vivo-expanded EPCs, highlighting a link between VP1u binding to the apical domain of hTfR and viral internalization. RBD mutants that disrupt its interaction with hTfR barely inhibited B19V infection in EPCs. These findings support a receptor-switch model in which AXL mediates attachment and hTfR drives internalization. Defining these mechanisms provides a foundation for developing antiviral strategies targeting B19V entry into EPCs.

## Introduction

Human parvovirus B19 (B19V) is a small, nonenveloped virus within the *Erythroparvovirus* genus of the *Parvoviridae* family (1). Its 5.6 kb single-stranded DNA (ssDNA) genome is flanked by identical inverted terminal repeats (ITRs) at both the 5′ and 3′ ends (2,3). B19V infection is globally prevalent and is the causative agent of fifth disease (erythema infectiosum) in children. In addition, B19V causes severe hematological disorders, including chronic anemia in immunocompromised individuals, nonimmune hydrops fetalis in pregnant women, and transient aplastic crisis in patients with increased erythrocyte hemolysis and high demand for increased erythrocyte production, such as those with sickle cell disease and hereditary spherocytosis (3–6).

During erythropoiesis, hematopoietic stem cells (HSCs) differentiate through defined stages, including burst-forming unit erythroid (BFU-E) progenitors, colony-forming unit erythroid (CFU-E) progenitors, and erythroblasts, ultimately giving rise to mature erythrocytes in the bone marrow and fetal liver (7). Erythropoiesis is tightly regulated by erythropoietin (Epo) within the hypoxic bone marrow environment (8,9). B19V preferentially infects early-stage erythroid progenitor cells (EPCs), including BFU-E and CFU-E progenitors, as well as pronormoblasts (10–18), and this selective tropism underlies its capacity to disrupt erythrocyte production and cause diseases (3,11,19). *Ex vivo*-expanded CD36⁺ EPCs differentiated from HSCs isolated from bone marrow are highly permissive to B19V infection and serve as a surrogate model of B19V infection, particularly when cultured under hypoxic conditions (20–22). In addition, UT7/Epo-S1 cells, a megakaryoblastoid leukemia cell line, can be infected by B19V and are commonly used for mechanistic studies (23).

The B19V capsid is ∼25 nm in diameter and exhibits T=1 icosahedral symmetry composed of 60 structural proteins, ∼5% VP1 (83 kDa) and 95% VP2 (58 kDa), which differ by a 227-amino-acid (aa) N-terminal extension unique to VP1, termed the VP1 unique region (VP1u) (24,25). The B19V VP1u is a distinguishing feature among parvoviruses. Unlike other parvoviruses, this region is highly immunogenic and a high number of neutralizing epitopes were found in aa 1-80 (26–31). B19V VP1u contains two key functional domains required for infection: the N-terminal region (aa 5–68), defined as the minimal receptor-binding domain (RBD), which mediates viral attachment and internalization (32,33), and a downstream phospholipase A2 (PLA_2_) domain (aa 128–160), which is required for escape from intracellular membrane vesicles (34). VP1u has been reported to be externalized on the capsid surface after capsid interaction with cells, prior to virus entry (35,36). Notably, recombinant VP1u or its RBD protein is sufficient to enter EPCs and other B19V-permissive cells but not B19V-nonpermissive cells, highlighting the central role of VP1u in B19V entry and cell tropism (32,33,37). Therefore, identification of the cellular entry co-receptor that interacts with B19V VP1u remains a central question in the field.

Through a genome-wide CRISPR-Cas9 guide RNA screen for host factors required for VP1u RBD entry, we identified that tyrosine-protein kinase receptor UFO (AXL) as a proteinaceous receptor for B19V infection of EPCs (33). B19V entry into EPCs is mediated by the interaction of VP1u RBD with host AXL expressed on B19V-permissive cells. Both the recombinant AXL ectodomain (ECD) and an anti-AXL polyclonal antibody block B19V infection of EPCs (33). Notably, the AXL tyrosine kinase activity is dispensable for B19V infection (33), suggesting that AXL primarily functions as an attachment receptor mediating B19V docking at the cell surface. However, how VP1u mediates B19V internalization and intracellular trafficking in EPCs remains incompletely defined.

In this study, we employed an engineered ascorbate peroxidase 2 (APEX2)-based proximity labeling coupled with quantitative mass spectrometry (qMS) to identify host proteins that interact with VP1u during viral internalization and intracellular trafficking (38–40). Using a recombinant VP1u-APEX2 fusion protein to capture host cell proteins within ∼20 nm of where VP1u localizes after internalization into UT7/Epo-S1 cells. This approach identified transferrin receptor 1 (hTfR) as a highly enriched protein. hTfR is a type II transmembrane glycoprotein that mediates iron uptake via clathrin-mediated endocytosis and is highly expressed in EPCs (41), where it plays a critical role in erythropoiesis. Its apical domain is known to engage multiple ligands, including ferritin heavy chain (FTH1) (42,43) and serves as an entry receptor for several viruses, including canine parvovirus (CPV) (44–50).

We further investigated the interaction between B19V VP1u and hTfR and defined the role of this interaction in viral internalization and infection. Our findings identify hTfR as a key entry co-receptor that mediates B19V internalization and may facilitate intracellular trafficking of the virus through direct engagement with the VP1u RBD.

## Results

### Identification of B19V VP1u-associated host proteins during VP1u entry into cells using APEX2-mediated proximity labelling

B19V VP1u is externalized on the capsid surface and mediates B19V virion entry into cells through direct binding to host cell receptors (32,33,37). To identify host cell proteins involved in B19V VP1u internalization and intracellular trafficking, we engineered a recombinant VP1u-APEX2 protein and an APEX2 control (**Figure 1A&B**). To assess whether the recombinant VP1u-APEX2 protein retains its capability to enter B19V-permissive UT7/Epo-S1 cells, we performed a protein internalization assay. The results showed that VP1u-APEX2 was internalized into the cells after incubation for 2 h at 37°C, as detected by anti-Flag, but not the APEX2 control (**Figure 1C**), confirming VP1u-dependent internalization of VP1u-APEX2.

**Figure 1.**
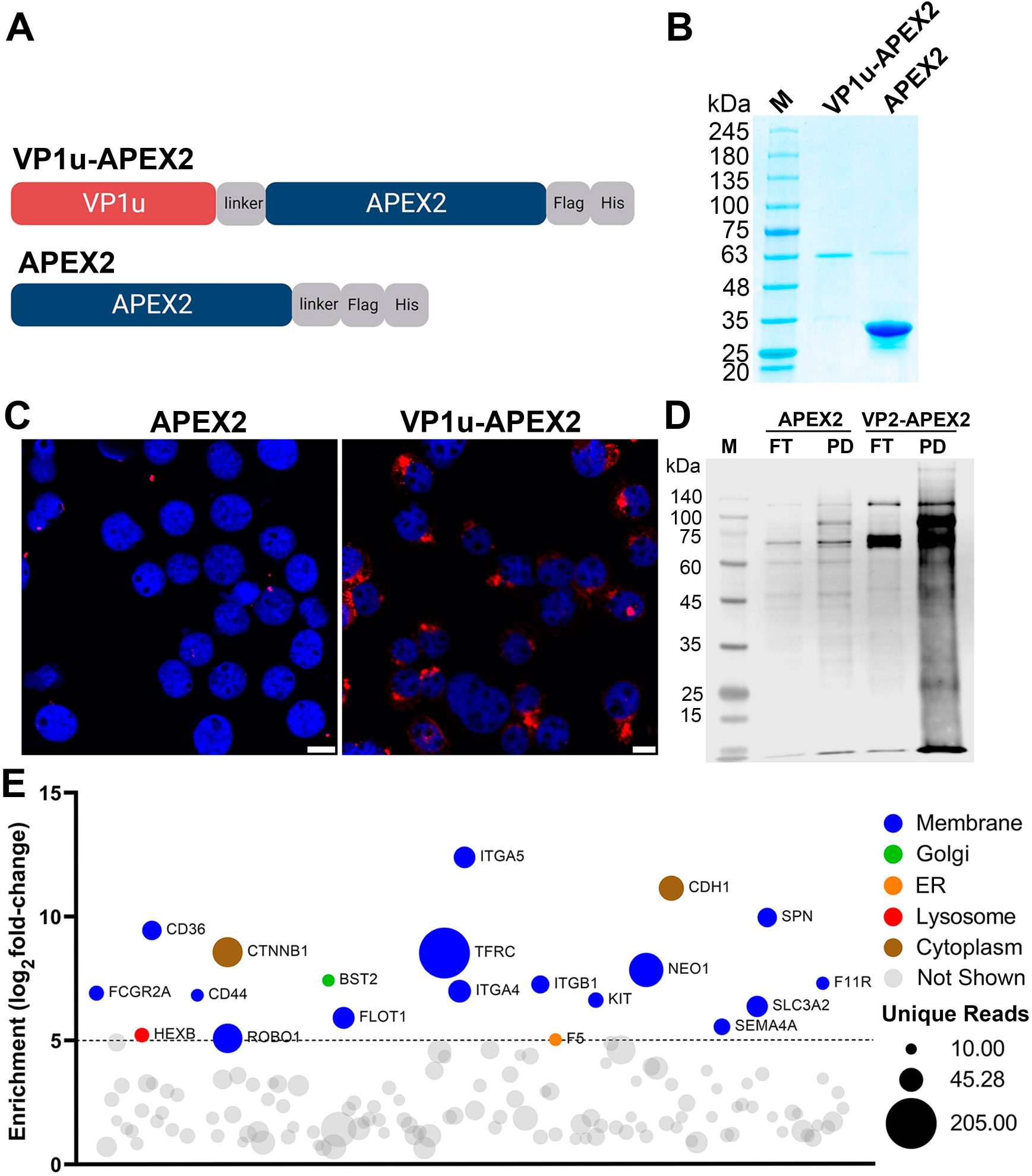
Identification of B19V VP1u-associated host proteins by APEX2-mediated proximity labeling. (A&B) VP1u-APEX2 and APEX2 control proteins. (A) Protein diagram. VP1u-APEX2 consists of APEX2 fused to the C-terminus of the unique region of B19V VP1 (VP1u) via a seven-residue glycine-serine linker (GGSGGSG), followed by a Flag tag and a 6 × Histidine (His) tag. APEX2 has a linker-Flag-His tag fused at the C-terminus. (B) Analysis of purified proteins. VP1u-APEX2 and APEX2 proteins were expressed in bacteria and purified. Approximately (∼) 1 µg of each protein was separated by SDS-PAGE, followed by Coomassie blue staining. M, molecular weight marker. **(C) Confocal microscopy of VP1u-APEX2 entry.** 1 × 10^6^ UT7/Epo-S1 cells were incubated with 2 μM VP1u-APEX2 or APEX2 protein at 37°C for 2 h. The cells were then immunostained with α-Flag to visualize internalized proteins under a Leica STED confocal microscope. Scale bar = 10 μm. Nuclei were stained with DAPI (4’,6-diamidino-2-phenylindole). **(D) Western blotting of APEX2-biotinylated proteins.** 1 × 10^7^ UT7/Epo-S1 cells were incubated with 2 μM VP1u-APEX2 or APEX2 protein at 37°C. After 2 h, APEX2-mediated biotinylation was then performed as described in the Materials and Methods and **Figure S1**. Biotinylated host proteins were purified with streptavidin-conjugated magnetic beads. The supernatant was collected as the flow-through (FT), and the beads were further washed several times and eluted as the pull-down (PD). Both FT and PD samples were analyzed by SDS-PAGE and immunoblotting using Alexa Fluor 680-conjugated streptavidin. **(E) Analysis of VP1u-APEX2-biotinylated/associated proteins using quantitative mass spectrometry (qMS).** Three independent PD samples prepared from VP1u-APEX2 and APEX2 (control) treated cells were analyzed by on-bead digestion and qMS. MS data were processed and analyzed as described in the Materials and Methods. The bubble plot shows protein enrichment (log_2_ fold change) in the VP1u-APEX2 group relative to the APEX control, with color indicating subcellular localization based on Gene Ontology (GO) annotation. TFRC denotes human transferrin receptor 1 (hTfR).

Following the optimized internalization conditions, we next evaluated APEX2-mediated proximity-biotinylated proteins. UT7/Epo-S1 cells were incubated with VP1u-APEX2 or APEX2 for 2 h at 37°C, followed by labeling with biotin-phenol and hydrogen peroxide for 1 min.

Biotinylated proteins were enriched using streptavidin magnetic beads, and pull-down (PD) and flowthrough (FT) fractions were analyzed by SDS-PAGE followed by immunoblotting for biotinylated proteins (**Figure 1D**). VP1u-APEX2 samples showed markedly higher biotinylation efficiency and a greater diversity of enriched proteins compared to APEX2 controls. Notably, distinct bands at ∼180 kDa and ∼90 kDa were observed exclusively in VP1u-APEX2 samples. These bands, along with a shared ∼130 kDa band present in both conditions were detected on a Coomassie blue-stained SDS-PAGE gel (**Figure S1**), and were excised and analyzed by mass spectrometry (**Table S1**). Human transferrin receptor 1 (TFRC or hTfR) was identified as the top hit based on unique peptide counts in the ∼180 and ∼90 kDa bands and the second-highest hit in the ∼130 kDa band of the VP1u-APEX2 samples, but was absent in the APEX2 controls (**Figure S1**).

To further define the VP1u-associated proteome, proximity labeling experiments were performed in triplicate (**Figure S2**), followed by on-bead digestion and quantitative mass spectrometry (qMS). A total of 150 proteins with ≥ 10 unique peptides were identified in VP1u-APEX2 samples (**Table S2**), which were plotted based on mass intensity fold-change enrichment relative to APEX2 controls (**Figure 1E**). Gene Ontology (GO) analysis categorized these proteins by cellular localization. Again, TFRC/hTfR exhibited the highest intensity in VP1u-APEX2 samples with a log₂ fold change of ∼9 compared to APEX2 samples, and had the greatest number of unique peptides (**Figure 1E**). Given its high expression in erythroid progenitor cells, regulation during erythropoiesis, and established role in clathrin-mediated endocytosis (41), hTfR emerged as a strong candidate for further investigation in B19V entry.

### hTfR colocalizes with B19V VP1u in UT7/Epo-S1 and CD36^+^ EPCs and with the B19V capsid in infected CD36^+^ EPCs

We next examined whether hTfR colocalizes with B19V VP1u during entry. To model VP1u-mediated uptake, we utilized a previously established GFP-tagged VP1u (GFP-VP1u) recombinant protein to mimic B19V entry (32,33). UT7/Epo-S1 cells were incubated with GFP-VP1u or GFP control protein at 4°C for 2 h to allow surface binding, followed by 37°C for 1 h min to permit internalization. Immunofluorescence analysis revealed clear colocalization of GFP-VP1u with hTfR [Manders’ overlap coefficient (MOC)=0.75], whereas no signal was detected in GFP-treated cells (**Figure 2A**). Similar colocalization of the VP1u RBD with hTfR was observed in UT7/Epo-S1 cells incubated with VP1u receptor-binding domain (RBD) (**Figure S3**).

**Figure 2.**
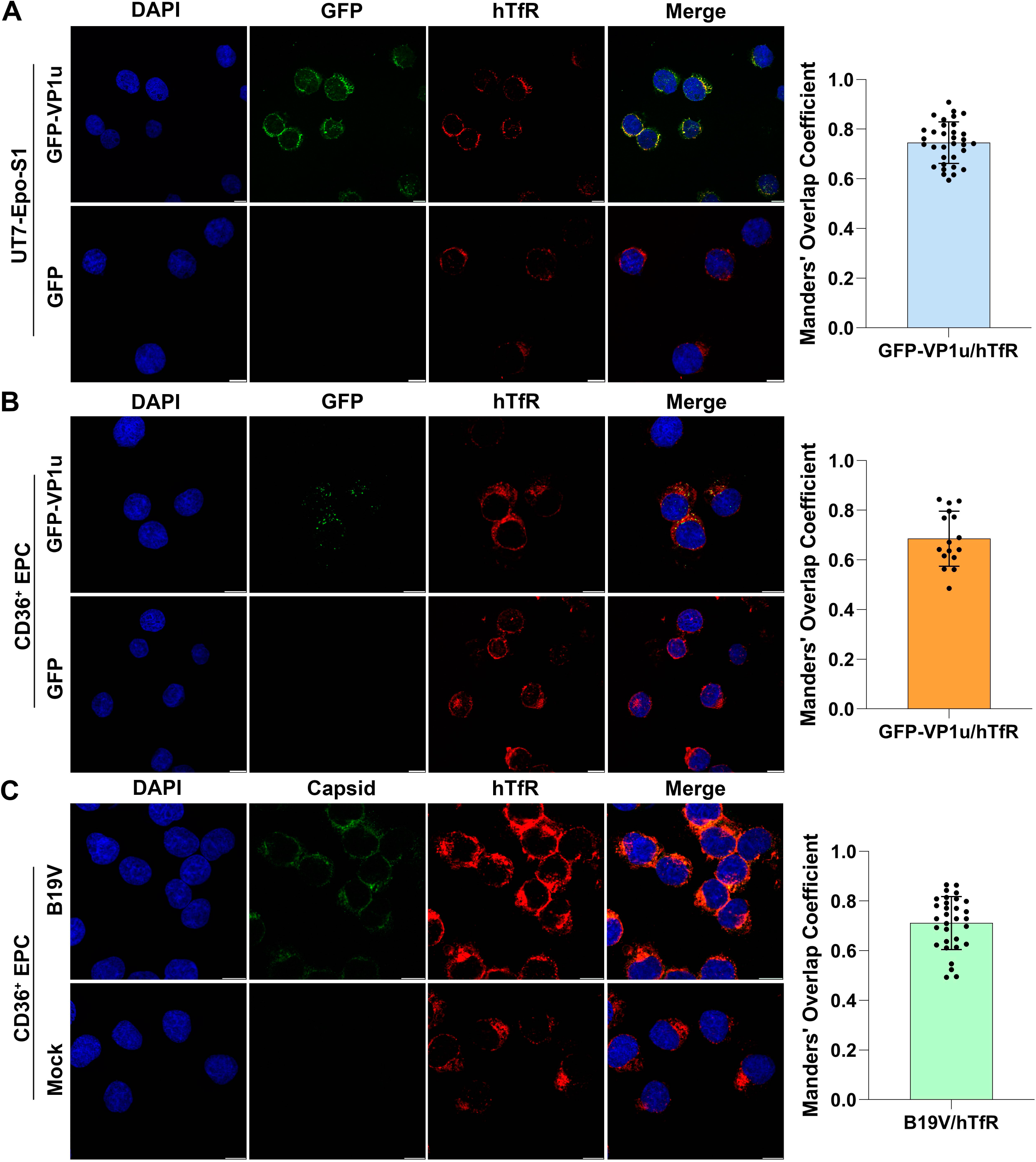
Human transferrin receptor 1 (hTfR) colocalizes with VP1u and B19V virions. (A&B) Colocalization of GFP-VP1u with hTfR. A total of 5 × 10^5^ UT7/Epo-S1 cells (A) or CD36^+^ EPCs (B) were incubated with 1 μM purified GFP-VP1u or GFP protein at 4°C for 2 h, followed by incubation at 37°C for 10 min. The cells were immunostained with α-hTfR (red). **(C) Colocalization of B19V capsids with hTfR in B19V-infected CD36^+^ EPCs.** A total of 5 × 10^5^ CD36^+^ EPCs were incubated with B19V at an MOI of 1 × 10^5^ vgc/cell at 37°C for 1h. The cells were immunostained with α-B19V capsid (green) and α-hTfR (red), with nuclei counterstained with DAPI (blue). Mock-infected cells served as a negative control. Scale bar = 10 µm. Colocalization was assessed using Manders’ overlap coefficient (MOC) for GFP-VP1u and hTfR (A&B) (S1, n ≥ 30; EPC, n = 16) and for B19V capsid and hTfR (C) in individual cells (n ≥30) using ImageJ (Fiji). Each dot represents one cell; bars indicate mean ± standard deviation (SD).

Furthermore, in primary CD36⁺ EPCs, a high level of colocalization between GFP-VP1u and hTfR was observed (MOC=0.69) (**Figure 2B**). To validate this finding in the context of infection, we assessed hTfR localization during entry of infectious B19V in CD36⁺ EPCs. Cells were infected at 37°C for 1 h and stained using a monoclonal antibody against the B19V capsid (clone 860-55D). Consistently, B19V capsid signals colocalized well with hTfR (MOC=0.71) (**Figure 2C**).

Together, these results demonstrate that both VP1u and infectious B19V virions colocalize with hTfR during the early stages of entry into B19V-permissive cells.

### hTfR is a critical host factor for B19V internalization and infection in UT7/Epo-S1 cells

To directly assess the functional role of hTfR in B19V infection, we next evaluated the internalization and replication of B19V in cells. Since CD36^+^ EPCs are highly dependent on transferrin to proliferate (21), we knocked down hTfR in UT7/Epo-S1 cells using an shhTfR RNA-expressing lentivirus (**Figure 3A**). We infected hTfR-knockdown cells (S1^shhTfR^) and scrambled shCtrl RNA-expressing lentivirus-transduced control cells (S1^shCtrl^) with B19V, and the internalized B19V genome was quantified. hTfR knockdown reduced B19V internalization to ∼49% of the control level (**Figure 3B**). We next assessed viral replication at 2 days post-infection (dpi). B19V DNA replication in S1^shhTfR^ cells was significantly decreased compared to S1^shCtrl^ cells (**Figure 3C**). To exclude the possibility that reduced replication resulted from global defects in host DNA synthesis on which B19V replication is dependent (51), we performed EdU incorporation assays in parental UT7-Epo/S1, S1^shCtrl^, and S1^shhTfR^ cells. hTfR knockdown did not alter host cell DNA synthesis or cell cycle progression in any of the three UT7-Epo/S1 cell types at 72 h post-transduction (**Figure 3D&E**), indicating that the observed reduction in viral replication is not attributed to impaired cellular proliferation.

**Figure 3.**
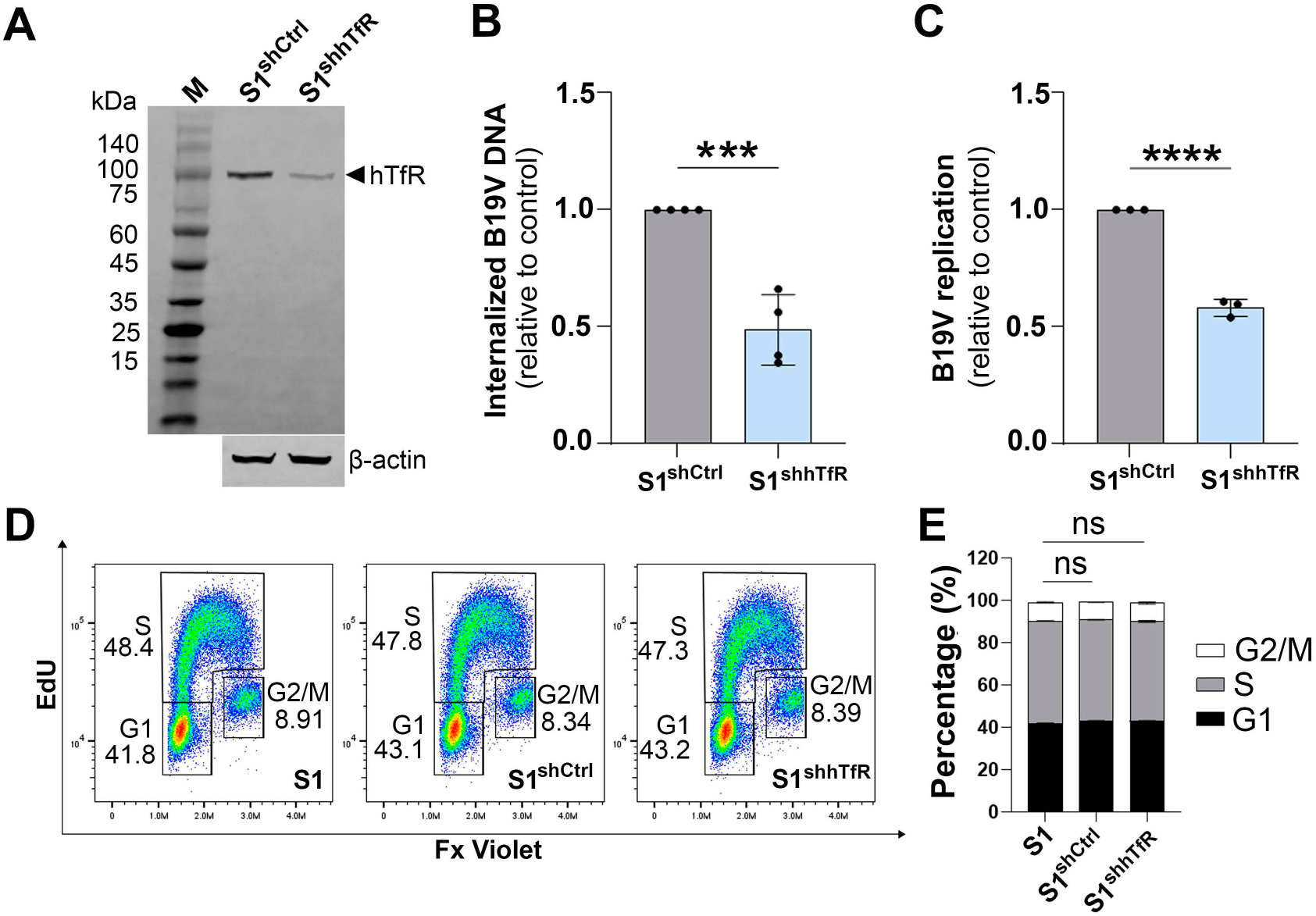
Knockdown of hTfR significantly reduces B19V internalization and infection in UT7/Epo-S1 cells but does not affect cell cycle progression. (A) hTfR knockdown (KD). UT7/Epo-S1 (S1) cells were transduced with shhTfR and shCtrl (scramble control)-expressing lentiviruses, respectively. S1^shhTfR^ and S1^shCtrl^ cells were lysed for Western blotting using α-hTfR (OKT9). β-actin serves as a loading control. **(B) Quantification of internalized B19V in hTfR-KD cells.** S1^shhTfR^ or S1^shCtrl^ cells were incubated with B19V at an MOI of 3,000 vgc/cell at 37°C for 1 h. After trypsin treatment, the cells were washed with PBS, and internalized viral DNA was extracted and quantified by measuring viral DNA levels normalized to mitochondrial DNA using multiplex qPCR. Relative values are shown. **(C) Quantification of B19V replication in hTfR-KD cells.** S1^shhTfR^ or S1^shCtrl^ cells were infected with B19V at an MOI of 1,000 vgc/cell. At 2 days post-infection (dpi), the cells were collected, and viral replication was quantified by measuring viral DNA levels normalized to mitochondrial DNA using multiplex qPCR. Relative values are shown. Data shown were analyzed by *t* test (\*\*\*\**P* < 0.0001). **(D&E) Analysis of cell cycle.** EdU incorporation assays were performed to monitor de novo DNA synthesis in the indicated cells, followed by flow cytometry analysis for cell cycle distribution. (D) Representative flow cytometry histograms showing cell cycle profiles of parental S1, control S1^shCtrl^ and S1^shhTfR^ cells. (E) Quantification of the percentage of cells in G2/M, S, or G1 phases. Data represent mean ± SD from three independent experiments. Statistical significance was determined by two-way ANOVA with Dunnett’s test (ns, not significant).

Together, these data establish hTfR as a critical host factor required for efficient B19V internalization and subsequent replication.

### B19V VP1u and VP1u RBD interact with hTfR in vitro with high affinity

We further assessed the direct interaction between hTfR and VP1u using a biolayer interferometry (BLI) assay. Using recombinant His-tagged hTfR ectodomain (hTfR^ECD^), the direct interaction with B19V VP1u was measured. hTfR^ECD^ was loaded onto NTA-coated biosensors at a concentration of 25 µg/mL for binding association and dissociation measurements with increasing concentrations of purified GST-VP1u and GST control (**Figure 4A**). A notable shift in the sensorgrams was observed with increasing concentration of GST-VP1u (**Figure 4B**). An equilibrium dissociation constant (K_D_) value of 128 nM was calculated from the K_a_ (association rate constant; 1/Ms) and K_d_ (dissociation rate constant; 1/s). Next, we assessed the direct binding between hTfR^ECD^ and VP1u^RBD^ under the same conditions. Again, we observed a dose-dependent shift in binding, with a K_D_ value of 86 nM (**Figure 4C**). As a control, GST did not show binding to hTfR (**Figure 4D**).

**Figure 4.**
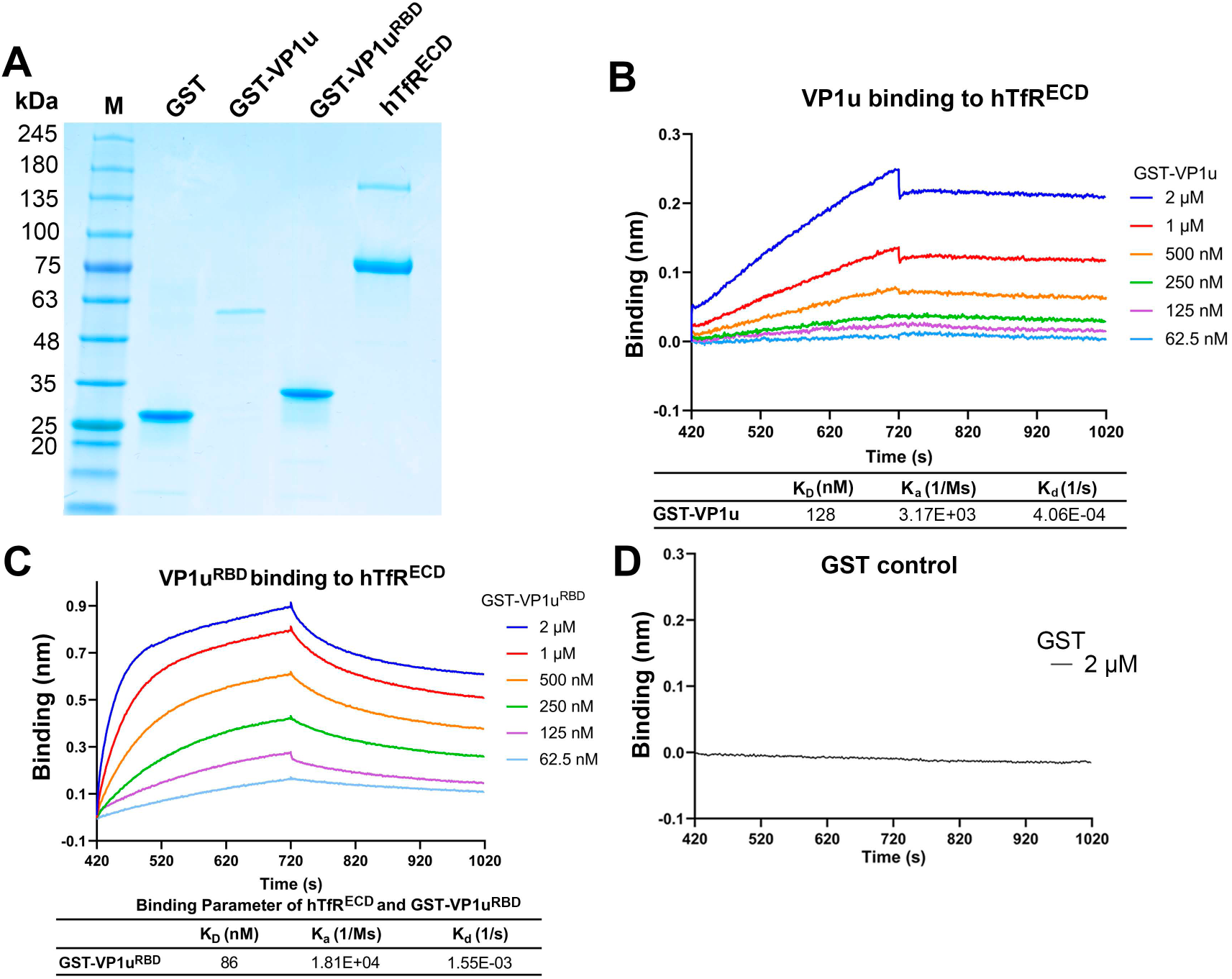
B19V VP1u and its receptor-binding domain (RBD) interact with hTfR ectodomain *in vitro*. (A) Expression and purification of recombinant proteins. GST, GST-VP1u, GST-VP1u^RBD^, and His-tagged hTfR ectodomain (hTfR^ECD^) proteins were analyzed by SDS-PAGE followed by Coomassie blue staining. A protein ladder was included. **(B-D) Binding kinetics of VP1u and VP1u^RBD^ to hTfR^ECD^ by biolayer interferometry (BLI).** Increased concentrations of GST-VP1u (B) or GST-VP1u^RBD^ (C) were tested for binding to hTfR^ECD^ (at 25 µg/mL) immobilized on NTA biosensors. Sensograms show dose-dependent association and dissociation. Kinetic parameters were derived by fitting the real-time binding curves, and the equilibrium dissociation constant (K_D_) was calculated as the ratio of dissociation rate constant [K_d_ (1/s)] and the association rate constant [K_a_ (1/Ms)]. 2 µM GST was used as a negative control to assess nonspecific binding (D).

Thus, these results indicate there is a direct, high-affinity interaction between hTfR and B19V VP1u.

### Blocking the hTfR apical domain with the monoclonal antibody OKT9 inhibits VP1u binding to hTfR, as well as B19V internalization and infection in EPCs

To directly test whether hTfR mediates B19V entry through its apical domain, we employed the mouse monoclonal antibody OKT9, which specifically targets the apical domain of hTfR (52), and has been shown to block viral engagement of hTfR (52–54). We first assessed whether OKT9 interferes with GST-VP1u^RBD^ binding to hTfR in vitro using BLI. Recombinant hTfR^ECD^ was immobilized on NTA biosensors and pre-incubated with OKT9 or isotype control IgG, followed by exposure to GST-VP1u^RBD^ (**Figure 5A**). Pre-binding of OKT9 markedly reduced GST-VP1u^RBD^ association with hTfR^ECD^, with dose-dependent inhibition (**Figure 5B**). We next evaluated the OKT9 inhibition in CD36⁺ EPCs. Cells pre-treated with OKT9 at 4°C exhibited substantially reduced internalization of GFP-VP1u following incubation at 37°C compared to IgG-treated controls, as quantified by confocal microscopy (**Figure 5C&D**). To determine the impact on B19V infection, CD36⁺ EPCs were pre-treated with OKT9. OKT9 treatment resulted in a strong, dose-dependent reduction in viral internalization (**Figure 5E**) and significantly impaired viral DNA replication at 2 dpi (**Figure 5F**). Importantly, OKT9 treatment did not affect host DNA synthesis or cell cycle progression (**Figure 5G&H**).

**Figure 5.**
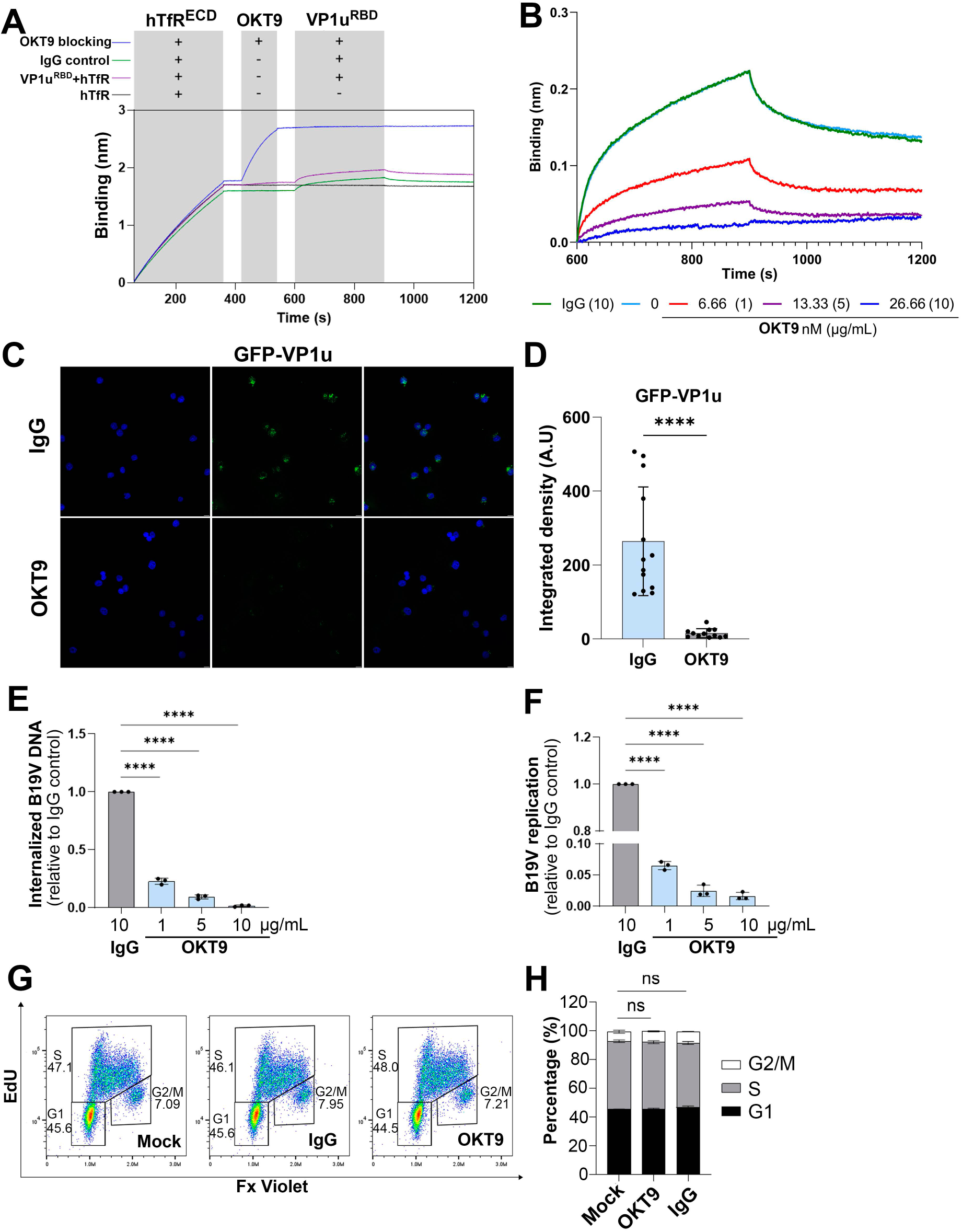
**hTfR monoclonal antibody, OKT9, blocks VP1u^RBD^-hTfR^ECD^ interaction and inhibits B19V internalization and replication in CD36^+^ EPCs, without affecting cell cycle progression. (A&B) BLI analysis of OKT9-mediated inhibition of VP1u^RBD^ binding to hTfR^ECD^**. (A) Experimental scheme. Binding of VP1u^RBD^ to hTfR^ECD^ immobilized on NTA biosensors was measured in the presence or absence of pre-incubation with the OKT9 blocking antibody or a mouse IgG control for non-specific binding. (B) Dose-dependent inhibition of VP1u^RBD^ binding to hTfR by OKT9. Sensograms show progressively reduced binding signals with increasing concentrations of OKT9, consistent with competitive inhibition of the VP1u^RBD^ to hTfR^ECD^ interaction. **(C&D) Immunofluorescence assay of OKT9 inhibition of VP1u entry into cells.** CD36^+^ EPCs were incubated with 10 µg/mL OKT9 antibody or mouse IgG control at 4°C followed by incubation with 1 μM GFP-VP1u at 37°C for 2 h. The cells were then fixed, permeabilized, and stained with DAPI, and were visualized for GFP signals for protein entry under a Leica STED confocal microscope (C). Scale bar = 10 µm. GFP intensity in cells was quantified using ImageJ (Fiji) in OKT9- and IgG-treated cell groups, respectively (D). Data shown are mean ± SD, and were analyzed by *t* test (**** P < 0.0001). **(E&F) Quantification of OKT9 inhibition of B19V internalization and replication.** (E) B19V internalization assay. CD36^+^ EPCs were pre-incubated with OKT9 or mouse IgG control at 4°C, followed by B19V infection (MOI=3,000) at 37°C for 1 h. After treatment with trypsin, the cells were washed with PBS, and viral DNA was extracted. Internalized B19V was quantified by measuring viral DNA levels normalized to mitochondrial DNA. (F) B19V replication assay. CD36^+^ EPCs were pre-incubated with OKT9 or mouse IgG control at 4°C followed by B19V infection (MOI=1,000) at 37°C. At 2 dpi, the cells were collected for quantification of B19V DNA levels. Data are presented relative to the corresponding control-treated cell group. Experiments were performed in triplicate, and data shown are mean ± SD, and were analyzed by one-way ANOVA with Dunnett’s test (****P < 0.0001). **(G&H) Cell cycle analysis following OKT9 treatment.** An EdU incorporation assay was performed in CD36^+^ EPCs under the indicated treatments, followed by flow cytometry analysis of cell cycle progression. (G) Flow cytometry histograms. Representative plots showing cell cycle profiles of untreated (mock), OKT9-treated (10 µg/mL), and IgG-treated CD36^+^ EPCs. (H) Quantification of the percentage of cells in G2/M, S, or G1 phase. Data represent the mean ± SD from three independent experiments. Statistical significance was determined by two-way ANOVA with Dunnett’s multiple comparisons test (****P < 0.0001; ns, not significant).

Together, these data demonstrate that OKT9 antibody-mediated blockade of the hTfR apical domain disrupts VP1u binding to hTfR and inhibits B19V internalization and infection in EPCs, supporting hTfR as a functional entry co-receptor.

### Ferritin heavy chain (FTH1) competes with VP1u for hTfR binding and inhibits B19V infection in EPCs

FTH1, an iron storage protein, is known to engage the apical domain of hTfR to mediate cellular entry and shares common hTfR contact regions with multiple pathogens (42,55). Based on this shared binding interface, we hypothesized that FTH1 may compete with VP1u for interaction with hTfR. To test this, we performed BLI to assess GST-VP1u^RBD^ binding to hTfR following FTH1 pre-binding. hTfR^ECD^ was immobilized on NTA biosensors and incubated with purified recombinant FTH1 (4 μM), followed by exposure to GST-VP1u^RBD^. Pre-binding of FTH1 inhibited GST-VP1u^RBD^ interaction with hTfR in a dose-dependent manner (**Figure 6A&B**). In contrast, maltose-binding protein (MBP) control showed no binding to hTfR and did not affect VP1u interaction with hTfR, confirming the specificity of FTH1-mediated inhibition. We next examined the functional consequences of this competition in CD36⁺ EPCs. Pre-treatment of CD36^+^ EPCs with FTH1 resulted in a strong reduction in B19V internalization (**Figure 6C**), and significantly decreased viral DNA replication (**Figure 6D**). Importantly, FTH1 treatment at concentrations up to 10 µM did not alter host DNA synthesis or cell cycle progression (**Figure 6E&F**).

**Figure 6.**
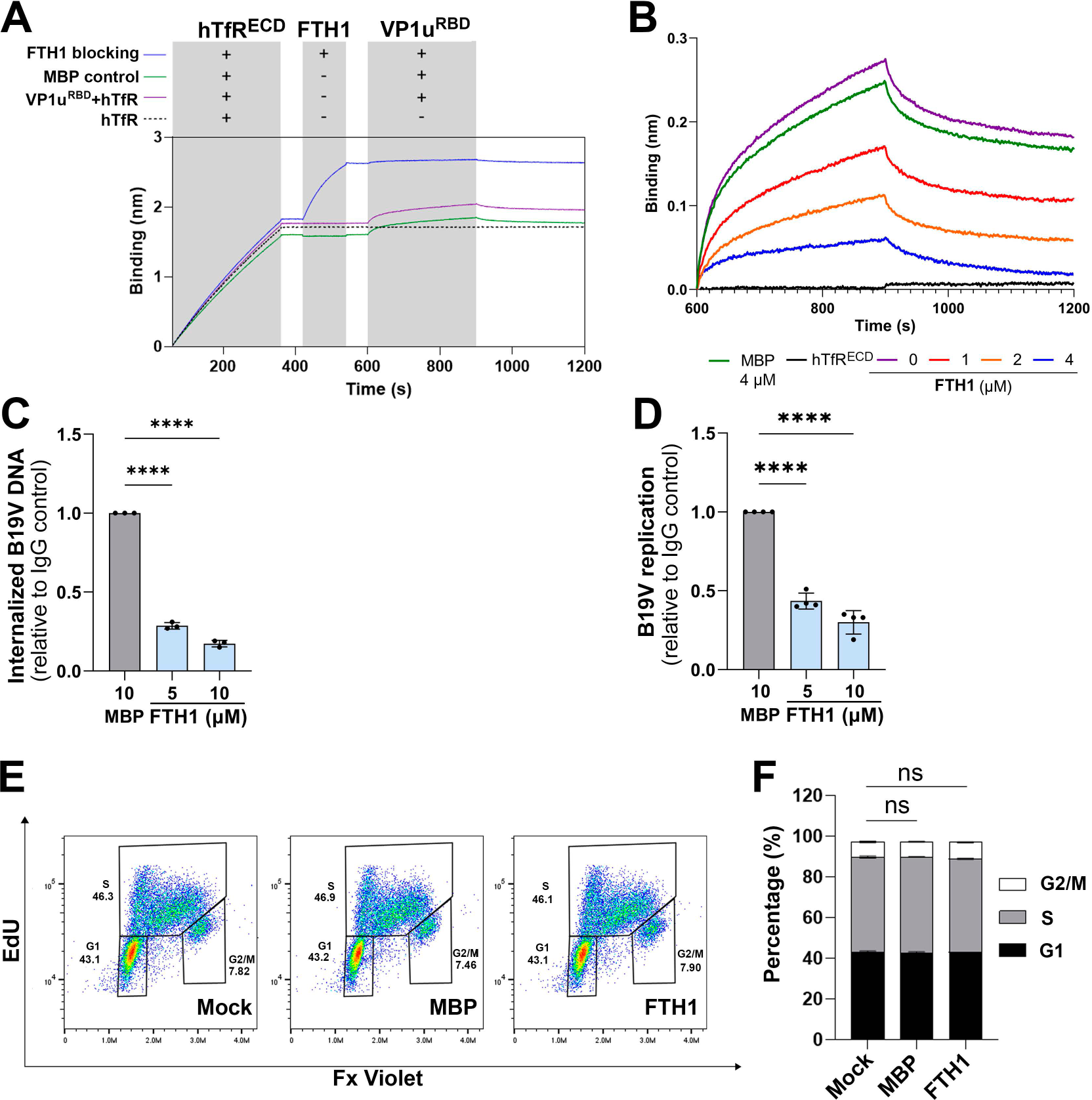
**Ferritin heavy chain 1 (FTH1) competes with B19V VP1u for binding to hTfR, thereby inhibiting B19V internalization and replication in CD36^+^ EPCs. (A&B) BLI analysis of FTH1 inhibition of GST-VP1u^RBD^ binding to hTfR^ECD^**. (A) Experimental scheme. hTfR^ECD^ was immobilized on Ni-NTA BLI biosensors, then introduced into the kinetic buffer with or without FTH1, and subsequently into buffer containing GST-VP1u^RBD^. Maltose-binding protein (MBP) served as a control for non-specific binding. (B) Dose-dependent inhibition of VP1u binding to hTfR by FTH1. Sensograms show progressively reduced association and dissociation of hTfR^ECD^ with GST-VP1u^RBD^ after the introduction of FTH1 at the indicated concentrations. MBP served as a control at 4 µM. **(C) Quantification of FTH1 inhibition of B19V internalization.** CD36^+^ EPCs were incubated with FTH1 or MBP control (at the indicated concentrations) at 4°C, followed by B19V infection (MOI=3,000) at 37°C for 1 h. After treatment with trypsin, the cells were washed with PBS, and viral DNA was extracted for quantification of internalized B19V by relative viral DNA levels normalized to mitochondrial DNA. **(D) FTH1 inhibits B19V infection.** CD36^+^ EPCs were pre-incubated with FTH1 or MBP control (at the indicated concentrations) at 4°C followed by B19V infection (MOI=1,000) at 37°C for 1 day. At 1 dpi, the cells were collected for quantification of B19V DNA levels. Data are presented relative to the corresponding control-treated cell group at each concentration. Experiments were performed in triplicate, and data shown are mean ± SD, and were analyzed by one-way ANOVA with Dunnett’s test (****P < 0.0001) **(E&F) Cell cycle analysis.** An EdU incorporation assay was performed in CD36^+^ EPCs with the indicated treatments, followed by flow cytometry analysis. (E) Flow cytometry histograms. Representative histograms show the cell cycles of mock, 10 µM FTH1-, and 10 µM MBP-treated CD36^+^ EPCs. (F) Quantification of the percentage of the indicated cell populations in G2/M, S, or G1 phase. Data shown are the mean ± SD from three independent experiments. Statistical significance was determined by two-way ANOVA with Dunnett’s multiple comparisons test (****P < 0.0001; ns, not significant).

Together, these findings demonstrate that FTH1 competitively inhibits VP1u-hTfR interaction and suppresses B19V internalization and infection in EPCs, further supporting the apical domain of hTfR as a critical interface for B19V VP1u engagement.

### B19V VP1u engages the apical domain of hTfR through defined structural determinants

We next sought to define the structural determinants within the VP1u RBD required for interaction with hTfR. Structural modeling using AlphaFold3 predicted a binding interface between VP1u RBD and the apical domain of hTfR (**Figure 7A**). Specifically, residues R208 and V210 of hTfR were predicted to interact with E54 within the loop between Helices 2 and 3 of VP1u RBD, while hTfR residue L212 was predicted to contact Q22 within Helix 1. To assess the importance of the VP1u RBD structure for binding to hTfR, we generated three VP1u RBD mutants designed to disrupt individual α-helices (RBDh1m, RBDh2m, and RBDh3m) (**Figure 7B**). Recombinant GST-tagged RBD and its mutant proteins were purified (**Figure 7C**). Binding analysis by BLI revealed that disruption of helix 1 or helix 3 (RBDh1m and RBDh3m) completely abolished binding to hTfR^ECD^, whereas the helix 2 mutant (RBDh2m) retained only residual binding (**Figure 7D**). Purified proteins were then used to inhibit B19V infection of CD36⁺ EPCs. The results show that while wild-type VP1u RBD effectively inhibited infection, all three mutants exhibited significantly reduced inhibitory activity (**Figure 7E**), consistent with the in vitro binding data.

**Figure 7.**
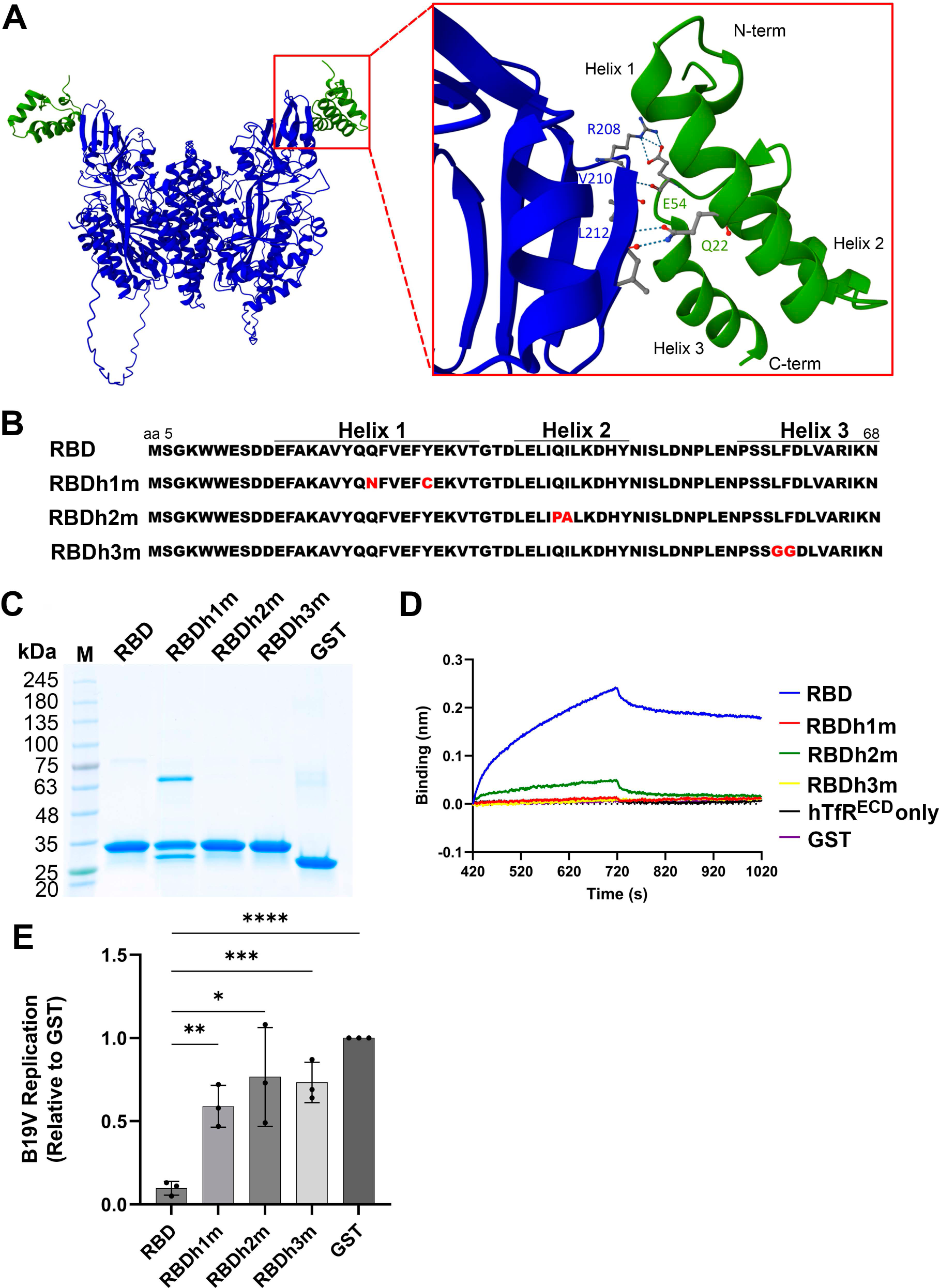
Structural modeling and mutational analysis of VP1u RBD required for hTfR binding and B19V infection in CD36^+^ EPCs. (A) Predicted structure model of the hTfR^ECD^–VP1u^RBD^ interaction. Structure complex predictions for hTfR^ECD^ and B19V VP1u^RBD^ were generated using the AlphaFold3 server, with the top-ranked model selected. The graphical representation of the model was created using Chimera, showing the VP1u RBD (green) engaging the apical domain of hTfR (blue). A magnified view highlights key interacting residues, including VP1u residue (e.g., E54 and Q22) and hTfR residues (e.g., R208, V210, and L212), suggesting a defined binding interface. **(B) Sequence and mutational design of VP1u^RBD^.** The RBD is organized into three predicted helices (Helices 1-3). Mutants (RBDh1m, RBDh2m, and RBDh3m) contain targeted substitutions highlighted in red. **(C) Purification of the mutated GST-VP1u^RBD^ proteins.** GST-tagged RBD and mutant proteins (RBDh1m, RBDh2m, and RBDh3m) were analyzed by SDS-PAGE with Coomassie blue staining. GST served as a control. M, protein size ladder. **(D) *In vitro* interaction between hTfR^ECD^ and mutant VP1u^RBD^ proteins.** BLI sensograms show the association and dissociation of 1 µM GST-VP1u^RBD^ (RBD) and its mutant proteins with hTfR^ECD^ immobilized on NTA biosensors. GST alone was used as a control to assess non-specific binding. **(D) Functional impact of RBD mutants on inhibition of B19V infection.** CD36^+^ EPCs were incubated with 0.5 µM of GST, GST-VP1u^RBD^ (RBD) or mutant proteins at 4°C, followed by infection with B19V (MOI=1,000) at 37°C. At 2 dpi, the cells were collected for quantification of B19V DNA levels. Experiments were performed in triplicate, and data shown are mean ± SD, and were analyzed by *t* test (\**P* < 0.1; \*\**P* < 0.01; and \*\*\**P* < 0.001).

Together, these results confirmed that all three α-helices within the VP1u RBD are critical for mediating RBD binding to the hTfR apical domain and are required for efficient inhibition of B19V infection by competition with B19V VP1u for binding.

## Discussion

In this study, we used an APEX2-based proximity labeling approach to unbiasedly identify hTfR as a co-receptor mediating B19V internalization and define the molecular basis of its interaction with the VP1u RBD. Using in vitro binding assays, viral internalization and infection assessments, and RBD structure-guided mutagenesis, we provide evidence that B19V engages the apical domain of hTfR to facilitate internalization of B19V virions into erythroid progenitors.

The reliance of B19V entry on hTfR is consistent with erythropoiesis that generates B19V tropic cell types, BFU-E and CFU-E progenitors and erythroblasts, where hTfR is abundantly expressed and regulated by STAT5 (56,57), an essential factor that activates B19V DNA replication (58). Transferrin is essential for erythropoiesis and is actively internalized by binding to hTfR via clathrin-mediated endocytosis (CDE) (56,59,60). The enrichment of hTfR during erythropoiesis partly supports B19V’s high tropism for erythroid progenitors.

Mechanistically, our data support a model in which B19V engages the apical domain of hTfR through VP1u RBD to initiate internalization. Notably, transferrin, the canonical ligand of hTfR, binds to the membrane-distal ectodomain of hTfR, involving the helical and protease-like domains (61), which is distinct from the apical domain that mediates binding of human ferritin, multiple pathogens (44,45,62,63), and the monoclonal antibody OKT9 (52). All of these observations support a model in which B19V engages hTfR independently of transferrin binding. Consistent with this, blocking the apical domain with the monoclonal antibody OKT9 and FTH1 significantly reduces VP1u binding, viral internalization, and downstream replication.

Ferritin, a physiological ligand of hTfR at the apical domain (55), is present in blood at a broad physiological range (∼18–350 µg/L), corresponding to ∼0.04–0.78 µM (assuming a ∼450 kDa holoferritin complex) (64). We observed that 10 μM ferritin heavy chain 1 (FTH1) (monomer; ≈0.42 µM holoferritin) reduced B19V infection in EPCs by ∼70%. Normal blood levels of ferritin (measured as serum ferritin) vary by sex and physiological state, and inflammation, liver disease, or iron overload (65). Elevated circulating ferritin has been reported as a marker of virus-induced inflammation (66–68), raising the possibility that increased ferritin levels may influence B19V infection in human bone marrow. Ferritin is a ubiquitous intracellular iron storage protein predominantly localized in the cytosol of cells of the reticuloendothelial system, including bone marrow macrophages, and under physiological conditions, free (soluble) ferritin in the bone marrow extracellular space is minimal (69,70). Therefore, our findings suggest potential clinical applications that B19V infection may be related to the serum ferritin level of infected individuals that is an indicator of fee ferritin in bone marrow (70), and that elevated ferritin or exogenous application of ferritin (e.g., FTH1) may help mitigate acute B19V infection.

Our mutagenesis studies further define the structural determinants of the VP1u interaction with the apical domain of hTfR. Disruption of α-helical elements within VP1u RBD largely abolishes binding to hTfR and impairs the ability of VP1u RBD to compete with binding of VP1u exposed from B19V virions during entry. These results establish a direct structure-function relationship between VP1u conformation and receptor engagement, supporting a model in which specific helical motifs mediate docking onto the hTfR apical domain. However, structural studies of the VP1u-hTfR complex and the VP1u-AXL complex will be critical to fully resolve the molecular interfaces of VP1u RBD with AXL and hTfR, which will explain the choice/switch of these two receptors used by B19V VP1u during entry.

Many viruses use TfR for entry through CDE (71). B19V utilizes VP1u to engage AXL and hTfR at the cell surface, triggering internalization via CDE (**Figure 8**). In this model, initial viral attachment is facilitated by AXL, an attachment receptor (33), rather than globside (Gb4), which was initially thought to be the primary attachment receptor for B19V (72,73), followed by high-affinity engagement of the hTfR apical domain by VP1u to drive uptake via CDE (**Figure 8**, Steps 1&2). This division of labor between attachment and entry receptors is a common feature of viral entry mechanisms, such as adenovirus, which uses CAR for attachment and integrin for entry (74,75), and may help explain the narrow tropism of B19V for EPCs (3,15).

**Figure 8.**
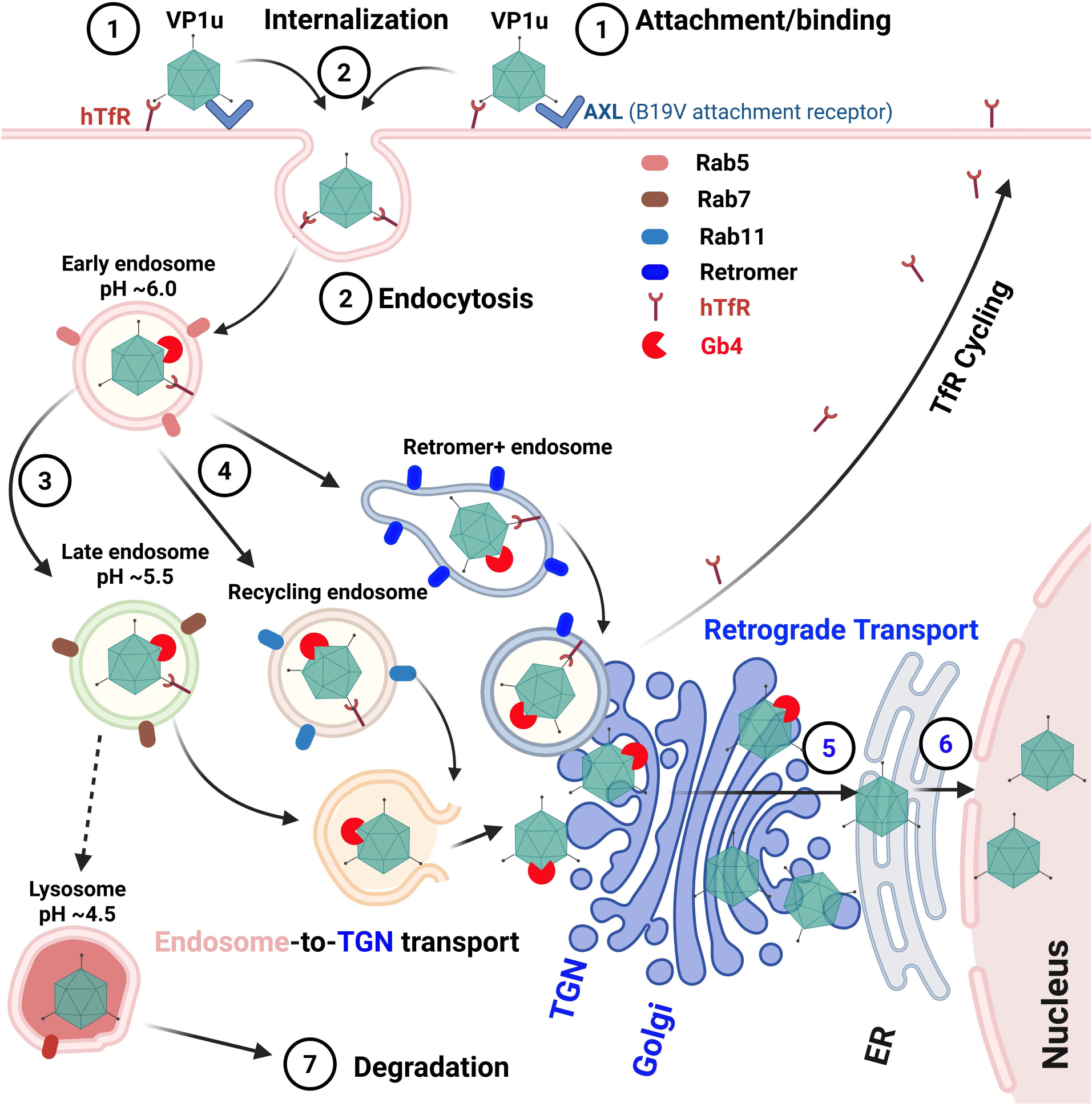
A model of parvovirus B19V binding and internalization (entry), intracellular trafficking, and productive infection. B19V initiates infection by binding to cell surface receptors through VP1u. (1) Viral attachment is mediated by AXL, which functions as an attachment receptor. (2) The virus is internalized via endocytosis, mediated by VP1u binding to hTfR. (3) hTfR facilitates B19V trafficking through early endosomes to late endosomes. A fraction of virions may undergo trans-Golgi network (TGN) localization. (4) B19V productive infection may involve other endosome-to–TGN transport pathways, including recycling endosome and retromer-facilitated retrograde transport. (5) Viral particles further follow retrograde trafficking to the Golgi and endoplasmic reticulum (ER). (6) From the ER or Golgi, they escape and gain access to the nuclear pore and enter the nucleus, where the viral DNA is uncoated, replicates, and is expressed. (7) Non-productive particles are trafficked to lysosomes for degradation. Globoside (Gb4) binds to B19V in endosomes and may escort them during retrograde transport. hTfR recycling occurs in parallel with viral trafficking.

Following internalization, the hTfR-transferrin complex formed at the cell surface is internalized by CDE and delivered to early endosomes (76), and recycling endosome, as well as potentially retromer-positive endosomes (**Figure 8**, Steps 3 and 4). Inside recycling endosomes, hTfR dissociates from transferrin and is recycled back to the cell surface (41,77). In these endosomes at an early stage, the low pH (∼pH 6.0-6.5) allows Gb4 to interact with B19V capsid, facilitating B19V intracellular trafficking toward the TGN and Golgi body (78,79). The virus then escapes from these membrane vesicles using the PLA_2_ domain within VP1u; however, PLA_2_ enzymatic activity requires elevated calcium concentration (Ca^2+^) and a neutral pH (80,81).

Notably, Ca^2+^ increases from ∼130 µM in TGN to >300 μM in the endoplasmic reticulum (ER) (82,83), while the pH rises from ∼6.0 in the TGN to ∼7.2 in ER (84,85). Thus, it is likely that B19V traffics through a retrograde pathway, via endosomes to the TGN and then to the ER, to penetrate the ER membrane for nuclear import (**Figure 8**, Steps 5&6). Nevertheless, the precise coordination between receptors, hTfR-mediated uptake and downstream trafficking factors warrants further investigation.

One of the essential questions in B19V infection is why progeny virus produced from B19V-infected EPCs is poorly infectious and the infectivity gradually declines (22). Therefore, infectivity studies are limited to using plasma samples containing high viral genome copy (vgc) levels of B19V, or virus purified from these samples. It is unclear whether VP1u is constitutively exposed on the virion surface or becomes accessible following initial receptor engagement, as previously suggested (35,36). We observed that B19V isolated from clinical specimens exhibits a broad range of VP1u exposure when analyzed by immunoblotting with an antibody targeting VP1u (**Figure S4**). Thus, we speculate that the degree of VP1u externalization on the capsid surface, in either clinical specimens or progeny virus produced from infected cells, is critical for conferring high infectivity.

In summary, our study identifies hTfR as a key entry co-receptor for B19V infection of EPCs and defines the molecular mechanism by which VP1u mediates viral internalization.

These findings position VP1u as a functional mimic or competitor of endogenous hTfR ligands, co-opting a native uptake pathway for viral entry, and thus as a critical determinant of B19V infectivity. By binding to the highly expressed AXL receptor, co-opting the hTfR endocytic pathway through direct engagement, and interacting with Gb4 in acidic endosomes, B19V exploits a central component of erythropoiesis to establish a productive infection. These findings provide a framework for understanding B19V entry and highlight potential targets for therapeutic intervention in clinical settings.

## Materials and Methods

### Primary cells and cell lines

**(i) Primary human CD36^+^ EPCs.** Primary human CD36^+^ EPCs were ex vivo expanded from CD34^+^ hematopoietic stem cells (HSCs), as previously described (22,58,86,87). Briefly, HSCs were cultured in Wong medium (21) under normoxia (5% CO_2_ and 21% O_2_) up to Day 4 (the day of procurement was designated as Day 0) and frozen in liquid nitrogen. In each experiment, Day 4 cells were seeded under normoxia for 2-3 days and transferred to hypoxic conditions (1% O_2_ and 5% CO_2_) for 2 days) (22), which was designated as Day 9 EPCs. These cells express erythroid cell markers CD36, EpoR (Epo receptor) and GPA (glycophorin A), as well as CD71 (hTfR) (56), and were designated as CD36^+^ erythroid progenitor cells (CD36^+^ EPCs).
**(ii) UT7/Epo-S1 cell line.** It is a clone of the human megakaryoblastoid UT7/Epo cell line, originally obtained from Dr. Kevin Brown at the Hematology Branch, NHLBI, NIH, with permission from Dr. Kazuo Sugamura (88). The UT7/Epo-S1 cells were cultured under normoxic conditions in Dulbecco’s modified Eagle’s medium (DMEM; #SH30022.01; Cytiva Life Science, Marlborough, MA) containing 10% fetal bovine serum (FBS; #F0926; MilliporeSigma, St. Louis, MO), 2 units/mL erythropoietin [Retacrit (epoetin alfa-epbx), Pfizer], 50 units/mL penicillin, and 50 µg/mL streptomycin.

### APEX2-mediated proximity labeling

APEX2-mediated proximity labeling was performed according to a published protocol with modifications (38,40). Briefly, 1 × 10^7^ UT7/Epo-S1 cells were incubated with 2 μM of purified VP1u-APEX2 or APEX2 protein at 37°C for 2 h with continuous gentle rotation. After 2 h, the cells were incubated with biotin-phenol (BP) (Biotinyl tyramide; #SML2135, MilliporeSigma) in culture media at a final concentration of 500 μM for 30 min at 37°C. Next, hydrogen peroxide (H_2_O_2_; #H1009, MilliporeSigma) was added at a final concentration of 1 mM for 1 min. The biotinylation reaction was then quenched by removing the media and washing cells using Quenching buffer containing the following quenchers: 10 mM sodium ascorbate (#S1349, Spectrum Chemical, New Brunswick, NJ), 5 mM Trolox (#238813, MilliporeSigma), and 10 mM sodium azide (#014314.22, ThermoFisher). After washing four times, the cells were centrifuged at 300 × g for 3 min. Pellets were lysed in radioimmunoprecipitation assay (RIPA) buffer (50 mM Tris-HCl, 150 mM NaCl, 0.1% SDS, 0.5% deoxycholate, 1% Triton X-100, pH 7.5) supplemented with quenchers, Protease Inhibitor Cocktail (PIC; #S8830, MilliporeSigma), and additional 1 mM phenylmethylsulfonyl fluoride (PMSF; #786–055, G-Biosciences, St. Louis, MO). Cells were incubated on ice for 2 min, and the lysate was clarified by centrifugation at 15,000 × g for 10 min at 4°C. The lysates were then incubated with streptavidin magnetic beads (#88817, Thermo Fisher) on a rotator for 1 h at room temperature. Next, the lysate-bead mixtures were collected and washed twice with RIPA, pH 7.5 (without quenchers, PIC, or PMSF), once with KCl (1 M), once with Na_2_CO_3_ (0.1 M), once with urea (2 M at pH 8.0), twice with RIPA (pH 8.0), and five times with TNS buffer (50 mM Tris-HCl, pH 8.0, 150 mM NaCl, 0.1% SDS). The washed beads (samples) were kept in TNS buffer and sent for quantitative mass spectrometry (qMS) at the Taplin Biological Mass Spectrometry Facility, Harvard University.

For sodium dodecyl sulfate-polyacrylamide gel electrophoresis (SDS-PAGE) or SDS-PAG followed by Western blot analysis of APEX2-mediated biotinylated samples, the lysate-bead mixtures were washed as described above and eluted in 3 x Laemmli buffer (187 mM Tris-Cl, pH 6.8, 4.5%SDS, 25% glycerol, 0.015% bromphenol blue, 5% DTT) supplemented with 2 mM biotin (#B4501, MilliporeSigma), and 20 mM dithiothreitol (DTT; #DTT25, GoldBio, St Louis, MO) for 5 min at 95°C. The supernatants were also collected and analyzed.

### On-bead digestion and liquid chromatography-tandem mass spectrometry (LC-MS/MS) analysis

The beads in TNS buffer were washed at least five times with 100 μL of 50 mM ammonium bicarbonate, then subjected to on-bead digestion with sequencing-grade trypsin (Promega, Madison, WI), and subjected to LC-MS/MS as described previously (40).

### Bioinformatic analysis of qMS data

The obtained qMS data were further analyzed based on unique peptide reads and peptide intensity (**Table S1**). Proteins with ≥ 10 unique peptides were analyzed to calculate p-values (Significance) using a *t* test (unpaired, two-tailed), and those with p > 0.05 were excluded. The log_2_ fold-change (FC) from the MS intensities of the detected peptides were calculated relative to the average intensity value of proteins with <10 unique reads. For functional classification, highly enriched proteins were further categorized using Gene Ontology (GO) analysis via PANTHER v19.0 (https://pantherdb.org/), and candidates were selected based on relevant functional pathways.

### Virus infection and quantification

**Virus infection:** Plasma samples containing B19V were provided by ViraCor Eurofins Laboratories (Lee’s Summit, MO). They were purified using CsCl gradient ultracentrifugation followed by dialysis against phosphate buffered saline (PBS, pH 7.4). The purified virus samples were quantified by quantitative (q)PCR using a B19V NS1-specific probe (89). CD36^+^ EPCs were infected with purified B19V at a multiplicity of infection (MOI) of 1,000 or 3,000 viral genome copies (vgc) per cell as indicated in the figure legends.

**Virus internalization assay:** For quantification of internalized virions, cells were infected with B19V at 37°C for 1 h and washed, followed by trypsin treatment (0.05% trypsin, 0.53 mM EDTA in PBS) to remove attached virus.

**Virus replication assay:** For quantification of viral DNA replication, at 2 dpi, viral DNA from the infected cells was extracted using the Quick-DNA/RNA Pathogen Kit (#R1043, Zymo Research Co., Irvine, CA). Extracted DNA was quantified using a NS1 probe-based qPCR for B19V DNA and mitochondrial DNA (Mito DNA) as previously described (90,91).

### Protein internalization assay

Protein internalization assay was performed as previously described (32,33). Briefly, 5 × 10^5^ UT7/Epo-S1 cells or CD36^+^ EPCs were harvested, washed cells were and resuspended in DPBS (#SH30028.03, Cytiva), pH7.4, at a final volume of 500 µL. The cells were then incubated with purified protein at 1-2 µM at 4°C for 2 h with continuous rotation, followed by incubation at 37°C for 10-30 min. After washing three times with PBS, the cells were spun onto slides using a Cytospin 4 (Thermo Scientific) for immunofluorescent assays. Nuclei were stained with DAPI (4’,6-diamidino-2-phenylindole).

### Lentivirus production and transduction

An hTfR-targeted small hairpin interference RNA (shhTfR) and a scrambled shRNA (shCtrl) were cloned into lentiviral vector pLKO-mCherrry (92). The sequence of shhTfR is 5’-GCA AAT GCT GAA AGC TTA AA T-3’. The shhTfR- and shCtrl-expressing lentiviruses were produced according to previously described methods (56,86). The transduction units of lentiviruses were titrated by qPCR Lentivirus Titer Kit (#LV900, Abm, BC, Canada) (93,94). UT7/Epo-S1 cells were transduced with the lentivirus at an MOI of ∼30 transduction units per cell.

### Antibody or recombinant protein inhibition of B19V infection

For antibody inhibition, OKT9 monoclonal antibody (mouse IgG), or a mouse IgG control was incubated with CD36^+^ EPCs at concentrations ranging from 1 to 10 µg/mL in DPBS for 1 h at 4°C. For protein inhibition, purified proteins, FTH1 or an MBP control, were incubated with CD36^+^ EPCs at various concentrations in DPBS for 1 h at 4°C. The cells were then infected with B19V, and viral replication was assessed by the quantification of viral DNA using qPCR.

### SDS-PAGE and Western blotting

For SDS-PAGE, purified proteins were dissolved in 1 x Laemmli buffer, heated at 95°C for 5 min, and separated on 4-20% pre-made gels (#4561095, Bio-Rad). The gels were stained with Coomassie blue.

For Western blotting, cells were collected and lysed as previously described (87). The lysates were separated by SDS-PAGE and transferred to a polyvinylidene difluoride (PVDF) membrane. The membrane was blocked with 5% non-fat milk in Tris-buffered saline buffer, pH7.4, and incubated with a primary antibody, followed by an infrared dye-conjugated IgG secondary antibody. Finally, the membrane was imaged on a LI-COR Odyssey F imager (LI-COR Biosciences, Lincoln, NE).

### Cell proliferation and cell viability assays

A 5-ethynyl-2’-deoxyuridine (EdU) incorporation assay was performed using the Click-iT™ Plus EdU Alexa Fluor™ 488 Flow Cytometry Assay Kit (#C10632, Invitrogen) to analyze the cell proliferation (cell cycle), followed by flow cytometry analysis using a 5-laser spectral flow cytometer (Aurora; Cytek Biosciences, Seattle, WA). FxCycle Violet (#F10347, Invitrogen) was used for flow cytometric analysis of DNA content in cells. Data were processed and analyzed using FlowJo v10 software (FlowJo LLC, Ashland, OR).

### Immunofluorescence assay and confocal microscopy

Cells were cytospun onto slides, fixed with 4% paraformaldehyde (PFA) for 15 min, permeabilized with 0.5% Triton X-100 for 5 min, and blocked with 5% fetal bovine serum (FBS) in PBS for 30 min. Cells were then incubated with primary antibodies followed by fluorophore-conjugated secondary antibodies. Nuclei were stained with DAPI. Images were captured using a confocal microscope (Leica TCS SP8 STED).

### Biolayer interferometry (BLI) assay

BLI was performed using an Octet RED96e (Sartorius, Bohemia, NY). NTA biosensors (#18-5101) were pre-equilibrated in kinetic buffer (25 mM Tris-HCl, 150 mM NaCl, 0.08% TWEEN20, pH 7.5) for 10 min before loading them with His-tagged hTfR^ECD^ protein at 25 μg/mL in kinetic buffer for 300 seconds (s) to allow proper binding. A baseline reading was recorded after protein loading in the kinetic buffer for 60 s to ensure stability. The biosensors were dipped into wells containing interacting proteins at varying concentrations for 300 s, and binding to hTfR^ECD^ was monitored in real-time. The biosensors were transferred back into the kinetic buffer, and the dissociation of interacting proteins from hTfR^ECD^ was recorded for 300 s. Raw BLI sensor data were analyzed using Octet Data Analysis Software for kinetic parameters: association rate constant (ka), dissociation rate constant (kd), and equilibrium dissociation constant (K_D_).

### Recombinant protein expression and purification

**Plasmid constructs:** VP1u-APEX2 or APEX2 encoding sequences as diagrammed in **Figure 1A** were synthesized at Twist Bioscience, South San Francisco, CA, and cloned into pET-30a(+) vector.

**Expression and purification:** Plasmids were transformed into *Escherichia coli* BL21(DE3) pLysS competent cells (Promega, Wisconsin, WI), and proteins were expressed by induction of 1 mM IPTG (isopropyl-β-D-thiogalactopyranoside). His-tagged proteins were purified using Ni-NTA agarose (Qiagen, Germantown, MD), according to the manufacturer’s instructions, or using Bio-Scale Mini Nuvia IMAC cartridges (#7800811, Bio-Rad) on a NGC Chromatography System (Bio-Rad).

Recombinant GST-VP1u, GST-VP1u^RBD^, GST, GFP-VP1u, or GFP-VP1u^RBD^ were expressed and purified as previously described (32).

### Antibodies and proteins used in the study

**Primary antibodies:** Mouse monoclonal anti-B19V capsid antibody (#MAB8292, clone 521-5D) was purchased from MilliporeSigma. Human monoclonal antibody against B19V conformational epitope (#Ab04448-10.0, clone 860-55D) was purchased from Absolute Antibody (Oxford, United Kingdom). Mouse monoclonal anti-hTfR antibody (#65605-1-MR, clone OKT9) was purchased from Proteintech. Rabbit polyclonal anti-hTfR antibody (#A5865) was purchased from ABclonal (Woburn, MA). Mouse anti-Flag antibody (#200-301-B13) was purchased from Rockland (Limerick, PA). Anti-B19V VP1u antibody was produced in-house (32).

**Secondary antibodies:** Alexa Fluor 488-conjugated goat anti-human IgG (H+L) secondary antibody, Alexa Fluor 555-conjugated donkey anti-mouse IgG (H+L) cross-absorbed secondary antibody (# A-32773), and Alexa Fluor 647-conjugated goat anti-rabbit IgG (H+L) cross-adsorbed secondary antibody (#A-21244) were purchased from Thermo Fisher. DyLight 800 conjugated anti-mouse IgG (#5257S) was sourced from Cell Signaling. Alexa Fluor 680-conjugated streptavidin antibody (#016-620-084) was purchased from Jackson Immunoresearch Laboratories Inc (West Grove, PA).

**Control antibodies:** Normal rabbit IgG control (#AB-105-C) were purchased from R&D Systems, Minneapolis, MN. Mouse IgG control (#SC2025) was purchased from Santa Cruz Biotechnology (Dallas, TX).

**Purified proteins:** Recombinant hTfR^ECD^ protein tagged with a His-tag at the C-terminus (#11020-H07H) and recombinant human ferritin heavy chain 1/FTH1 (#13217-HNAE) were purchased from SinoBiological (Paoli, PA). Recombinant Maltose-binding protein (MBP) (#NBC1-18538) was purchased from Novus Biologicals (Centennial, CO).

### Ethics statement

B19V-containing plasma samples were obtained from ViraCor Eurofins Laboratories (Lenexa, KS). Primary human CD34^+^ HSCs were isolated from the bone marrow of healthy human donors, and were purchased from AllCells LLC (Alameda, CA). Both the cell and virus samples were deidentified, and therefore institutional review board (IRB) review was waived.

### Data and materials availability

All data and materials used to evaluate the conclusions in this study are presented in the paper and/or the supplemental material. qMS data are available at Mass Spectrometry Interactive Virtual Environment (MassIVE) under accession number MSV000101293 (doi:10.25345/C5FX74C03).

### Statistics analysis

Statistical analysis was performed using GraphPad Prism version 9.5 and error bars represent the mean ± standard deviation (SD). Statistical significance P values were determined using t test, or one-, or two-way ANOVA. ****, P<0.0001, ***, P<0.001, **, P<0.01 and *, P<0.05 were regarded as statistically significant, and n.s. indicates not significant.

## Competing interest statement

S.M. and J.Q. have filed a provisional patent, no. 63/843,662 (Human Transferrin Receptor 1 as a Receptor Target for Parvovirus B19 Infection) on July 14, 2025.

## Acknowledgments

We are indebted to Dr. Richard Hastings and Marie Mahnke at the Flow Cytometry Core Laboratory of the University of Kansas Medical Center. We thank members of the Qiu lab for helpful discussions. The study was supported by National Institutes of Health grants AI130613; AI166293, AI180416, AI182645, AI150877, and HL174593. The funder had no role in study design, data collection and interpretation, or the decision to submit this work for publication.

---

**Figure S1.**
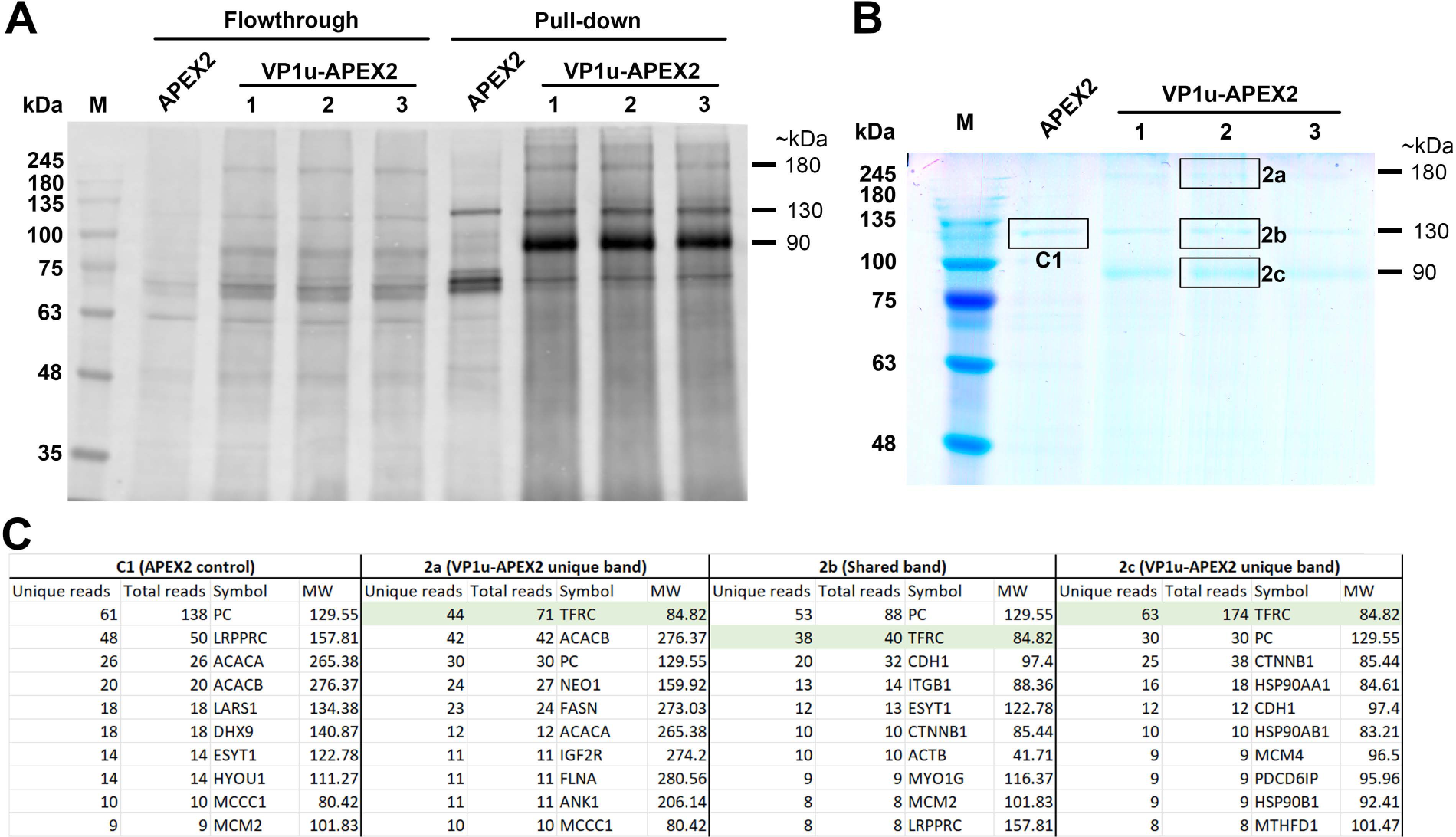
Identification of VP1u-APEX2 proximately biotinylated host proteins by SDS-PAGE and mass spectrometry. (A&B) Streptavidin pulldown of biotinylated proteins. A total of 1×10^7^ UT7/Epo-S1 cells were incubated with 2 μM of VP1u-APEX2 or APEX2 control proteins at 37°C for 2 h. APEX2-mediated proximity biotinylation was then performed as described in the Materials and Methods and **Figure S2**. (A) Immunoblotting for biotinylated proteins. Both flow-through (FT) and pulldown (PD, bead-bound) samples were analyzed by SDS-PAGE followed by immunoblotting using Alexa Fluor 680-conjugated streptavidin. (B) Coomassie blue staining biotinylated proteins. Pulldown samples were resolved by SDS-PAGE and visualized by Coomassie blue staining to assess protein enrichment and to guide excision of bands (2a, 2b, and 2c) and a control band (0b), as indicated, for downstream mass spectrometry. **(C) Mass spectrometry identification of proteins from excised gel bands.** Tables list the top 10 proteins identified from each band based on unique peptide counts, with corresponding molecular weights (MW) indicated. TFRC (hTfR) was identified as the most enriched protein in Bands 2a and 2c, corresponding to monomer (∼90 kDa) or dimer (∼180 kDa) forms, respectively, and ranked second in band 2b (∼130 kDa). The intermediate molecular weight of Band 2b likely reflects a differentially glycosylated form of hTfR. Additional proteins identified include cytoskeletal, chaperone, and membrane-associated factors.

**Figure S2.**
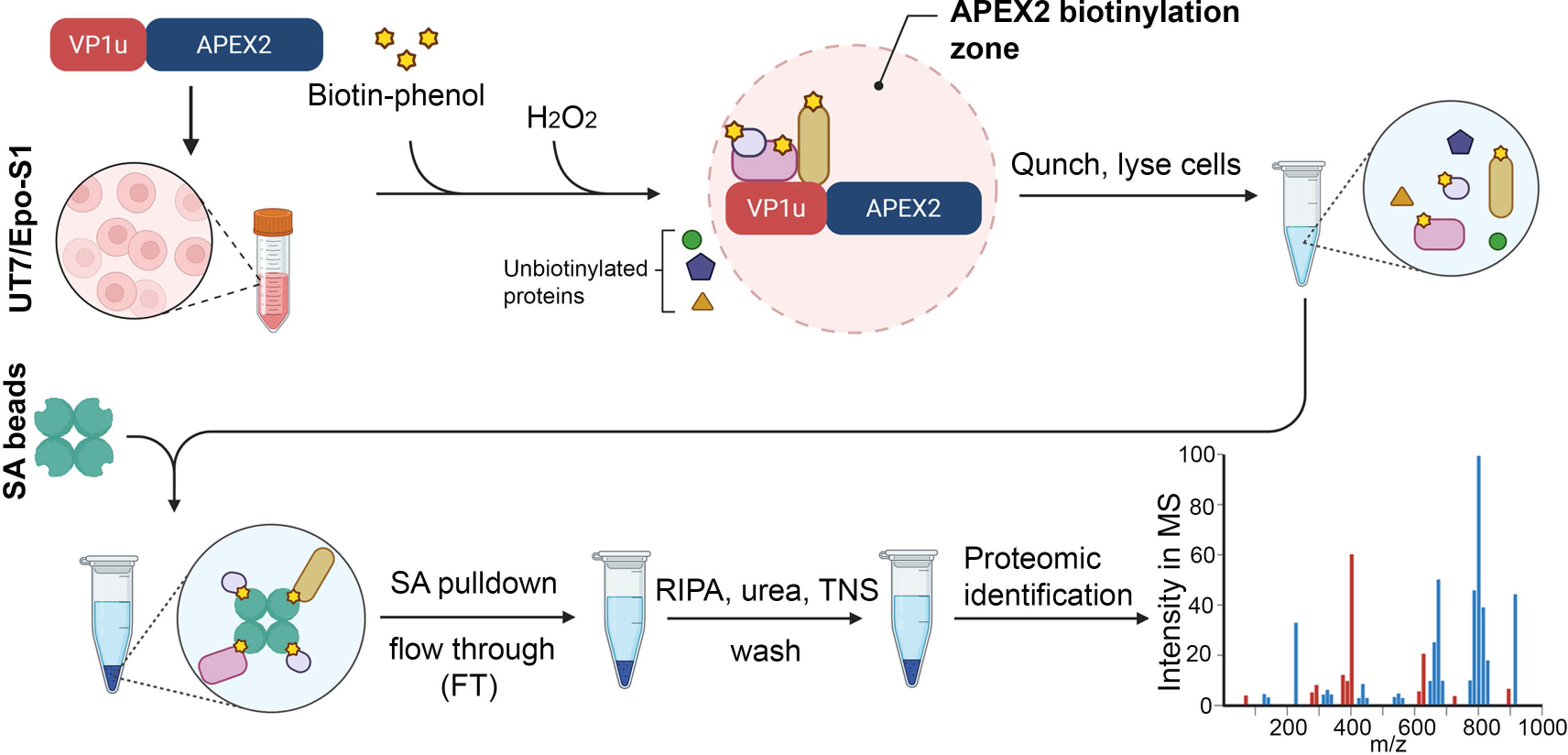
Diagram of workflow for APEX2-mediated proximity labelling. UT7/Epo-S1 cells were incubated with VP1u-APEX2 or APEX2 control in the presence of biotin-phenol (at 37°C for 30 min), followed by addition of H₂O₂ (1 min at room temperature) to induce biotinylation of proximal proteins. The reaction was then quenched, and the cells were lysed in RIPA buffer. The biotinylated proteins were enriched using streptavidin (SA) beads. After stringent washing (with RIPA, urea, and TNS buffers), bound proteins were subjected to on-bead enzyme digestion and proteomic identification by mass spectrometry.

**Figure S3.**
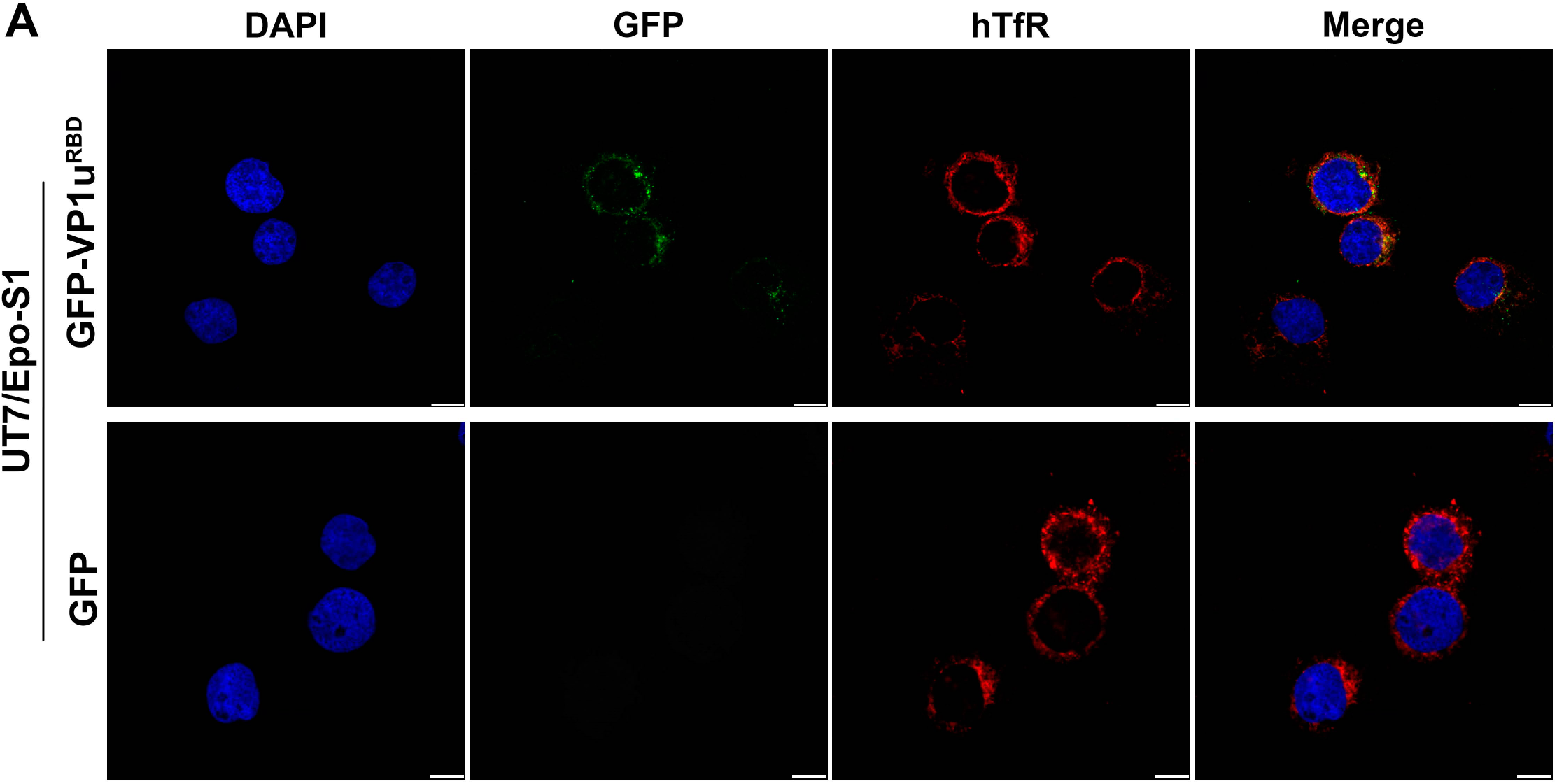
VP1u RBD colocalizes with hTfR in UT7/Epo-S1 cells. A total of 5 × 10^5^ UT7/Epo-S1 cells were incubated with 1 μM purified GFP-VP1u^RBD^ or GFP protein at 4°C for 2 h with continuous rotation, followed by incubation at 37°C for 10 min. The cells were then fixed, permeabilized, and stained with α-hTfR antibody. Nuclei were stained with DAPI. Bar = 10 µm.

**Figure S4.**
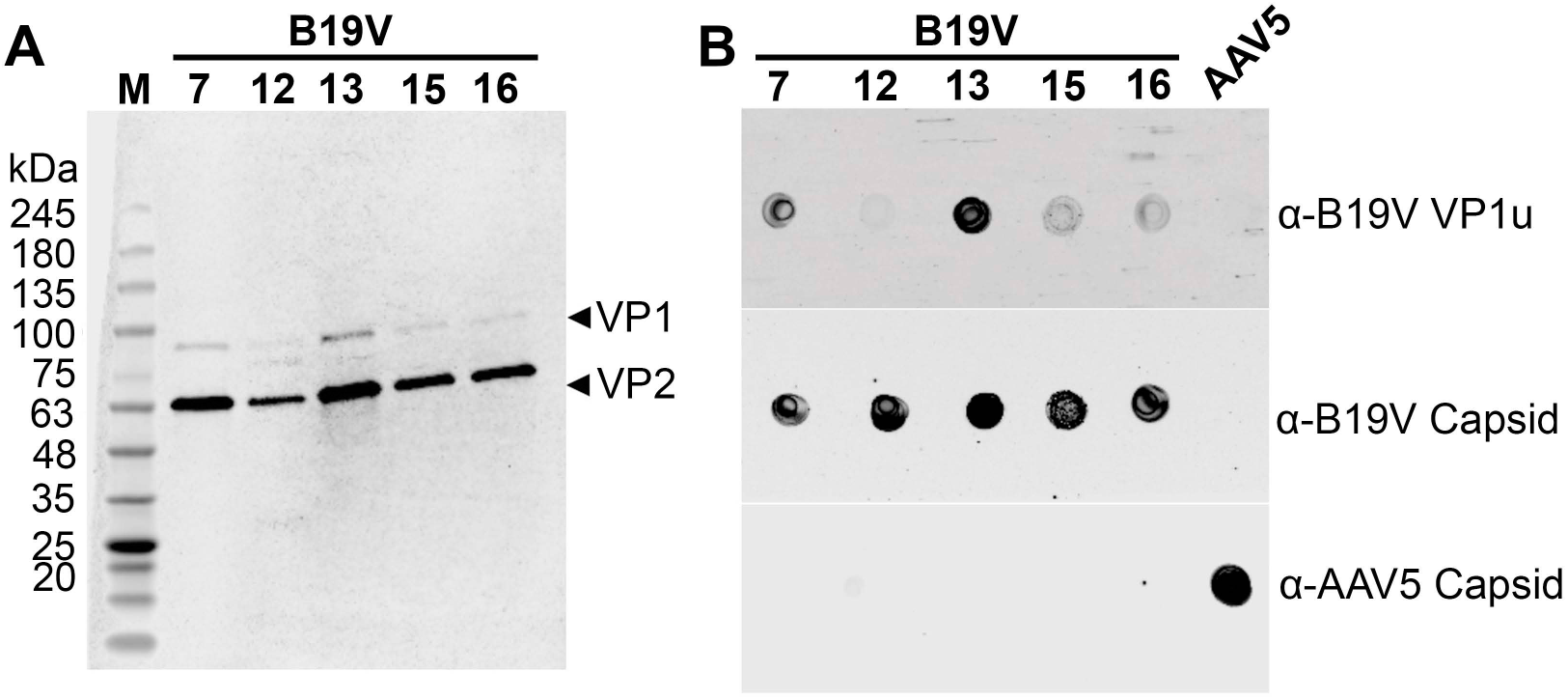
Characterization of purified B19V samples by Western blotting and dot blotting. (A) Western blot analysis of purified B19V. B19V plasma samples (#7, 12, 13, 15, and 16) were purified in CsCl equilibrium ultracentrifugation. Approximately 1 × 10^9^ viral genome copies (vgc) per sample were resolved by SDS–PAGE and probed with antibodies against B19V capsid proteins VP1 and VP2, as indicated. M, molecular weight marker (kDa). **(B) Dot blot analysis of B19V preparations** (**95**). Equal amounts (5 × 10^8^ vgc) of each sample were spotted onto nitrocellulose membranes (Amersham) and probed with antibodies against B19V VP1u (α-B19V VP1u) and B19V capsid proteins (α-B19V capsid; Clone 521-5D (33,96)). AAV5 was included as a specificity control and detected using an anti-AAV5 capsid antibody (Clone ADK5b).

**Table S1. Proteins identified from excised SDS-PAGE gel bands by mass spectrometry.**

**Table S2. Proteins identified by label-free quantitative mass spectrometry (qMS).**

